# Estimates of community stability using the invasion criterion are robust across levels of invader species richness

**DOI:** 10.1101/2024.04.18.590097

**Authors:** Meaghan Castledine, Daniel Padfield, Angus Buckling

**Affiliations:** College of Life and Environmental Sciences, Environment and Sustainability Institute, University of Exeter, Penryn, Cornwall, TR10 9EZ, U.K

**Author notes:** **Corresponding author:** Meaghan Castledine. **Author contributions:** MC and AB conceived and designed the study. MC conducted the experiments. MC and DP analysed the data. All authors contributed to the writing of the manuscript. **Competing Interests:** We have no competing interests. **Data availability:** All data and R codes used in the analysis will be available on GitHub.

## Abstract

A key feature of natural communities is that the species within them stably coexist. A common metric used to test community stability is measuring the ability of each species to recover from rare. Here, each species is assumed to have negative frequency dependent fitness and have a greater fitness relative to the other community members. A conceptual issue with measurements of relative invader fitness is that single species are invaded from rare. In natural communities, multiple species would likely decline following perpetuations e.g. antibiotic application, global warming, natural disasters. In our study, we compare previous estimates of community stability in a five species microbial community to experimental results in which multiple species are invaded from rare. Our results showed that single species invasions were broadly predictive of whole community stability when multiple species are invaded simultaneously. Precise values of relative invader fitness were less comparable, however being non-significantly different in most comparisons in 3/5 species. This work provides the first experimental test of the robustness of relative invader fitness metrics under multi-species invasion scenarios.

## Introduction

A key feature of natural communities is that the species within them stably coexist (Grainger *et al*. 2019). Stable coexistence can arise via multiple mechanisms such as differential resource use (Buche *et al*. 2022; Godoy *et al*. 2017; Li *et al*. 2019; Rainey & Travisano 1998; Schoener 1974; Stomp *et al*. 2004), higher-order interactions (Singh & Baruah 2021) and density-dependent predation (Coblentz & DeLong 2020; Gendron 1987; Winter *et al*. 2010). In theory, coexistence can be measured by a species’ ability to recover from rare following perturbation (Chesson 2000, 2018; Germain *et al*. 2018; Grainger *et al*. 2019; Siepielski & McPeek 2010). Assuming species have niches distinct enough for coexistence, a rare species should display negative frequency dependent fitness while resident species, greater in density, are limited by intraspecific competition (Chesson 2000, 2018; Grainger *et al*. 2019). The invasion criterion, which tracks the relative growth rate of rare populations compared to residents, has become a widely-used metric in community ecology (Grainger *et al*. 2019).

In modelling frameworks and experimental tests of mutual invasibility, single species are decreased to low densities while all other species are at high or equilibrium density (Grainger *et al*. 2019). A conceptual issue with this assumption is that perturbations are likely to affect more than one species simultaneously resulting in more than one species decreasing while others may be unaffected (Petraitis *et al*. 1989; Schwartz *et al*. 2020). If coexistence arises due to differential resource use, how many species become rare is unlikely to affect coexistence as each species is similarly limited by intraspecific competition (Chesson 2018). However, if mutual invasibility relies on higher-order or intransitive interactions among resident species, multiple species becoming rare may result in a resident species (now free from interactions that limited its impact on other species) competitively excluding an invader (Godoy *et al*. 2017; Singh & Baruah 2021). When networks follow intransitive dynamics, niche differences are required to promote coexistence (Godoy *et al*. 2017), suggesting instability if multiple species become rare. Similarly, if invasion is dependent on facilitation by another resident species, both species decreasing to low densities may result in loss of one or both species (Altieri *et al*. 2010; Stachowicz 2001). Understanding the validity of these metrics when multiple species recover from rare is important considering natural communities are exposed to stressors affecting multiple species simultaneously. Global warming, for instance, is introducing extreme weather events resulting in multiple populations declining (Sala *et al*. 2000). On smaller scales, microbiomes are broadly perturbed following antibiotic applications (Schwartz *et al*. 2020). Populations have the opportunity to recover when conditions become favourable again and whether recovery is successful depends on interactions between residents that may have increased in density and co-invaders (Petraitis *et al*. 1989).

In long term studies that follow natural communities, multiple species’ densities can fluctuate in response to changing environmental conditions (e.g. seasonal changes to temperature or resources) with differential responses from each species to each change facilitating coexistence (Adler *et al*. 2006; Angert *et al*. 2009; Descamps-Julien & Gonzalez 2005). For instance, two prairie grass species recovered from rare against one dominant species owing to variations in annual environmental conditions (Adler *et al*. 2006). Nevertheless, whether these species could invade from rare in a stable environment following brief perturbation is less understood as stable coexistence depends on predictable environmental fluctuations.

Consequently, we aim to address whether estimates of relative invader fitness (RIF) taken from single species invasions predict coexistence compared to scenarios in which multiple species invade from rare in a stable environment. Measurements of relative invasion ability have been predominantly conducted in microbes and plants as growth rates and generation times allow for relatively rapid assessment of coexistence and characterisation of interspecies interactions (Adler *et al*. 2006; Angert *et al*. 2009; Castledine *et al*. 2020b; Chesson 2018; Grainger *et al*. 2019; Rainey & Travisano 1998). Concurrently, we use a five-species microbial community consisting of soil bacteria which has been shown previously to be stable (Castledine *et al*. 2020b, a). We previously used estimates of single invasion RIF to predict the community’s stability by decomposing the community into all two to five species combinations (Castledine *et al*. 2020b). While these estimates support that all five species can stably coexist, can they when multiple species invade from rare within the five-species community? We make direct comparisons between assays in which resident species are the same (e.g. *Achromobacter* and *Pseudomonas*) and compare relative invader fitness for a focal species (e.g. *Variovorax*) in the absence (original assay) or presence (this study; e.g. *Stenotrophomonas* and *Ochrobactrum*) of co-invaders. We use these comparisons to determine if previous estimates of single-species RIF can predict the stability of the whole community (five species) when multiple species invade from rare.

## Methods

Bacteria species were originally isolated from soil and include: *Achromobacter* sp. (A), *Ochrobactrum* sp. (O), *Pseudomonas* sp. (P), *Stenotrophomonas* sp. (S) and *Variovorax* sp. (V) (Castledine *et al*. 2020b). Species can be identified by their unique colony morphology when plated on King’s medium B (KB) agar (Castledine *et al*. 2020b). In previous work, the stability of this community was characterised by constructing every possible combination of species, from two to five species richness. Within each combination, single focal species were invaded from rare (100-fold lower than resident species) (Figure 1). A community combination was defined as stable if the focal species’ growth rate compared to the resident species growth rates (relative fitness) was greater than 1. In most combinations the focal species had relative fitness values >1 indicating this community to be highly stable (71/75 significantly >1 and all combinations with means >1) (Castledine *et al*. 2020b).

**Figure 1.**
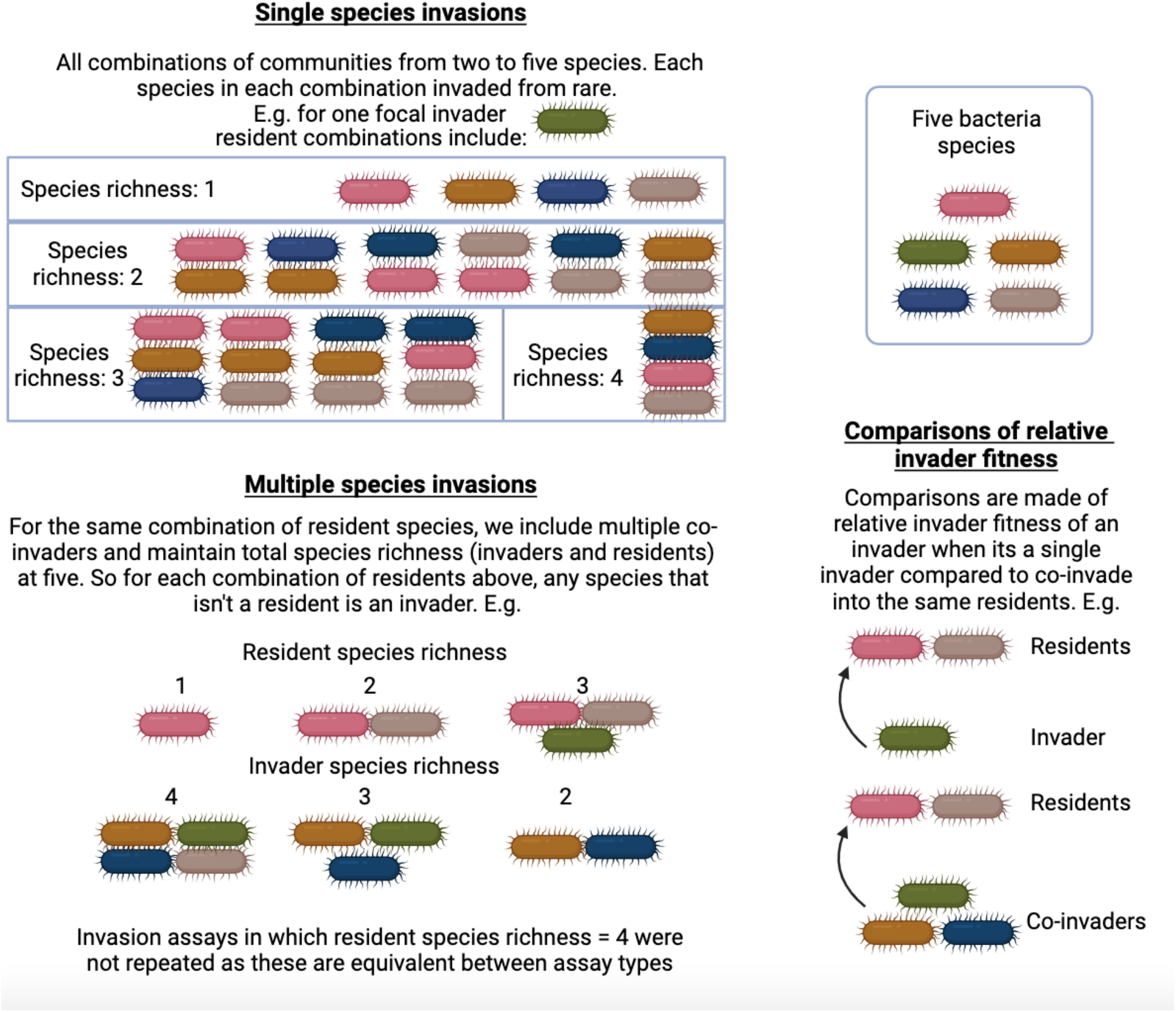
Species combinations for single invasion and co-invader assays. In previous work, single species are invaded into all possible combinations of resident species from one to four resident species richness. In this study, the same combinations of residents are used with any of the five species that is not a resident being an invading species. For example, when *Achromobacter* and *Ochrobactrum* are residents, *Stenotrophomonas, Variovorax* and *Pseudomonas* are invaders. As such, the relative invader fitness of each of these invaders can be compared to when they are invading as single invaders into communities of *Achromobacter* and *Ochrobactrum*.

**Figure 2.**
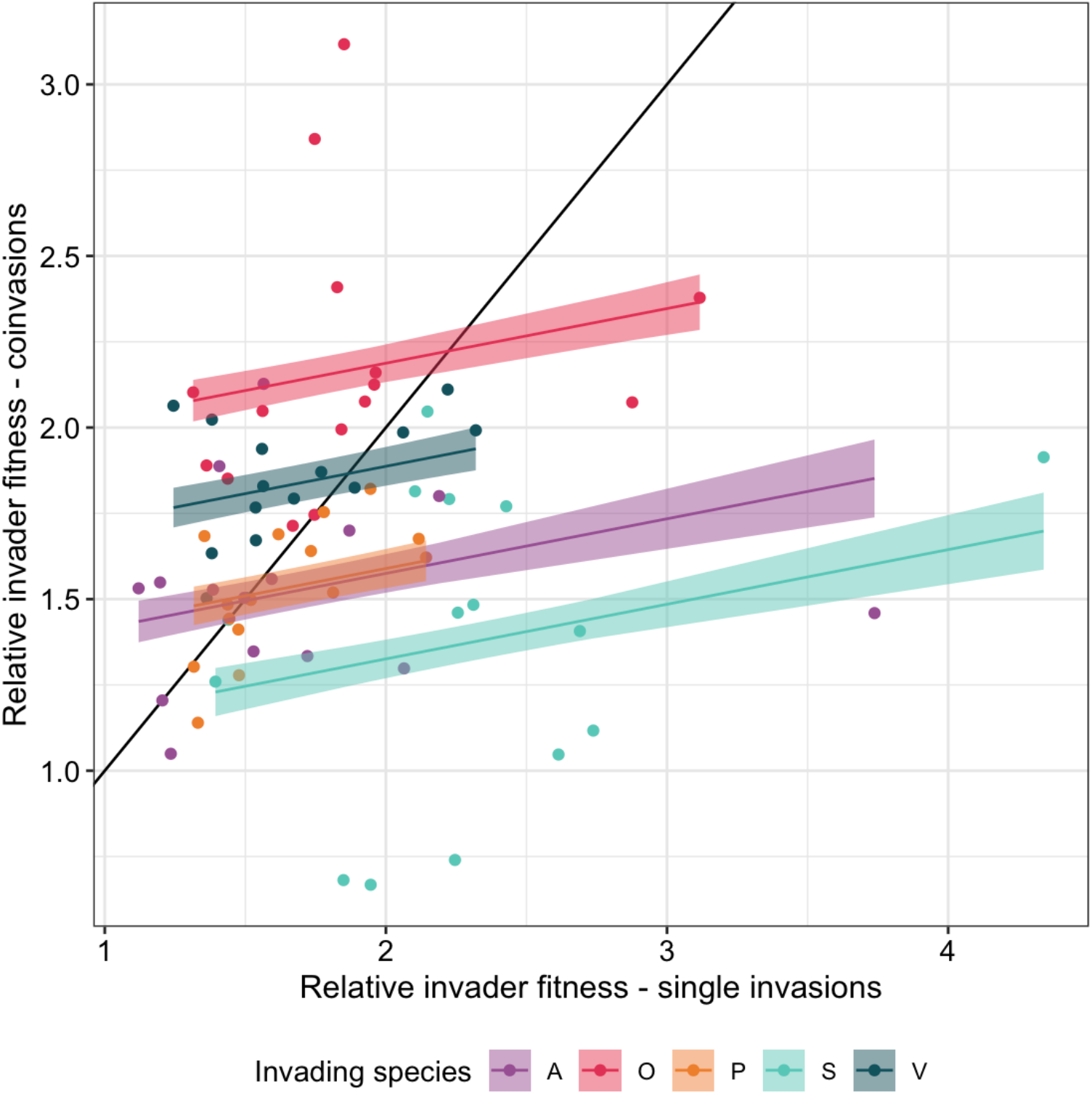
The relationship between relative invader fitness when each species is invaded as a single species compared to the presence of co-invaders. Lines and bands represent the best fit line and 95% confidence intervals of the model. The 1-1 line indicates the slope if estimates of relative invader fitness were predictable in the presence and absence of co-invaders.

**Figure 3.**
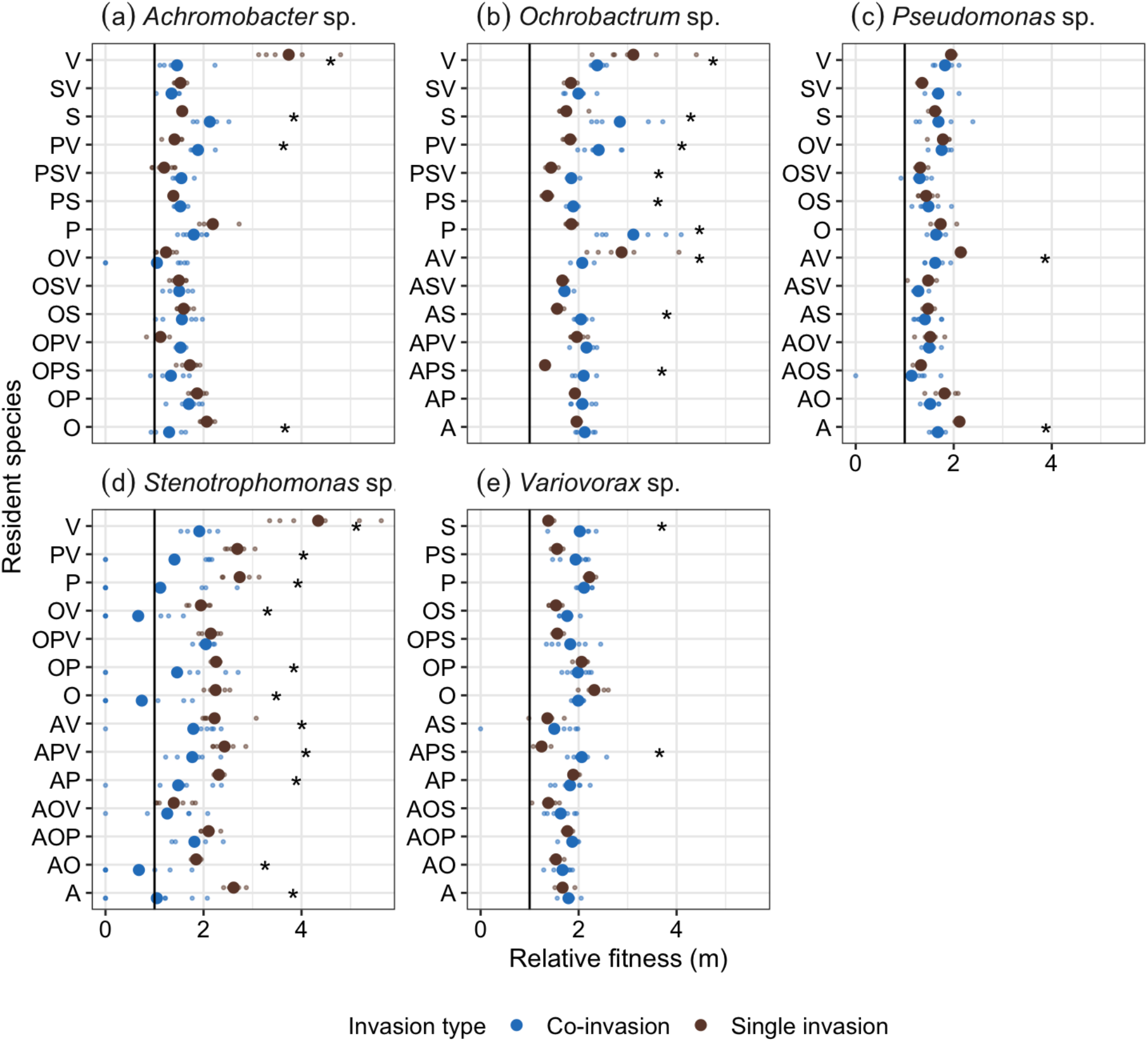
Comparisons of relative invader fitness with focal species (a-e) invading into different combinations of resident species in the presence or absence or co-invaders. Large points indicate mean values while small points indicate values from independent treatment replicates. X = 1 is the threshold where relative invader fitness values above this value indicates a stable community. Asterisk’s show significant differences in relative invader fitness estimates between invasion types. Resident species combinations are a combination of the first letter of each resident species (e.g. OP = *Ochrobactrum* and *Pseudomonas*).

In this study, community stability was tested by maintaining total species richness at five (all species always present) and instead manipulating which species were invading from rare from two to four invader richness (Figure 1). Single species invasions (one invader, four residents) were not conducted again, having been conducted in the aforementioned study. This resulted in a total of 70 treatments, each replicated six times. Any species which was not a resident was instead an invading species. Isogenic populations were grown in 6 mL 1/64 TSB (tryptone soy broth diluted with demineralised H_2_O) for two days, shaking (180 rpm) at 28°C in 25 mL glass vials with loosened plastic lids. Cell density (colony-forming units (CFU)) of each species was normalised to 10^5^ CFU/μL (method described in (Castledine *et al*. 2020b)). The sum of invading species were inoculated at a 100-fold lower density (1.6 μL, 16 x 10^4^ CFUs total) than the sum of the resident community (160 μL, 16 x 10^6^ CFUs total). Each species was represented at equal ratios within each invading or resident community. Communities were cultured in fresh 1/64 TSB, static at 28°C. After one week, samples were cryogenically frozen (−80°C) in glycerol (900 μL of culture with 900 μL 50% glycerol). Samples were plated from frozen onto KB (Kings medium B) agar and incubated for two days at 28°C. Relative invader growth rate of the individual invading species was calculated as the ratio of estimated Malthusian parameters (*m*), *m*_focal_:*m*_community_ where *m*_community_ is the total density change of all the other populations combined including residents and co-invaders and *m*_focal_ is the density change of one of the invading species. *m* = ln(N1 / N0)/t where N1 is the final density, N0 is starting density, and *t* is the assay time (1 week) (Lenski *et al*. 1991).

### Statistical analysis

All data were analysed using R v4.2.1 in R Studio and all plots for experimental data were made using the package ‘ggplot2’ (Team 2013; Wickham 2016). Model simplification was conducted using likelihood ratio tests and Tukey’s *post hoc* multiple-comparison tests were used to identify the most parsimonious model using the R package ‘emmeans’ (Lenth 2018).

First, we assessed whether relative invader fitness estimates under co-invasions significantly differed from 1 (indicating community stability). We performed independent one-sample t-tests on each combination of focal species by resident community combination, with the null-hypothesis being that the mean relative invader growth rate equals 1. A relative invader growth rate above 1 would indicate that the invader had a faster growth rate than the resident community, demonstrating negative frequency dependence. This resulted in 70 statistical tests and p-values were adjusted using the false discovery rate (*fdr*) method (Benjamini & Hochberg 1995).

Next, we assessed whether estimates of relative invader fitness correlated between single-species and co-invasions. The mean relative invader fitness of each species within each combination of resident community was taken in both datasets. In a linear model, mean relative fitness under co-invasion was analysed against interacting fixed effects of mean relative fitness under single invasion and species identity (which species the relative fitness metrics referred to). Next, comparisons were made of relative invader fitness (co-invading and single invasion treatments) within each combination of resident species. Here, relative fitness was analysed against interacting fixed effects of treatment (focal invading species into different resident species combinations) and invasion type (co-invasion or single invasion).

## Results

### The community is stable in the majority of co-invasion combinations

Contrary to previous estimates of community stability that estimate the ability of single species to recover from rare, we approached this by invading multiple species from rare within a five species community. In this study, 56/70 combinations were stable with relative invader fitness values significantly greater than 1 (Table 1; Figure 1). In 10/14 of the failed invasions, *Stenotrophomonas* was the invading species with populations being below detectable density in at least one replicate of each combination. In contrast, *Achromobacter* accounted for 2 failed invasions, and *Pseudomonas* and *Variovorax* 1 failed invasion each. In these three species, mean relative invader fitness was greater, but not significantly different, to 1.

**Table 1.**
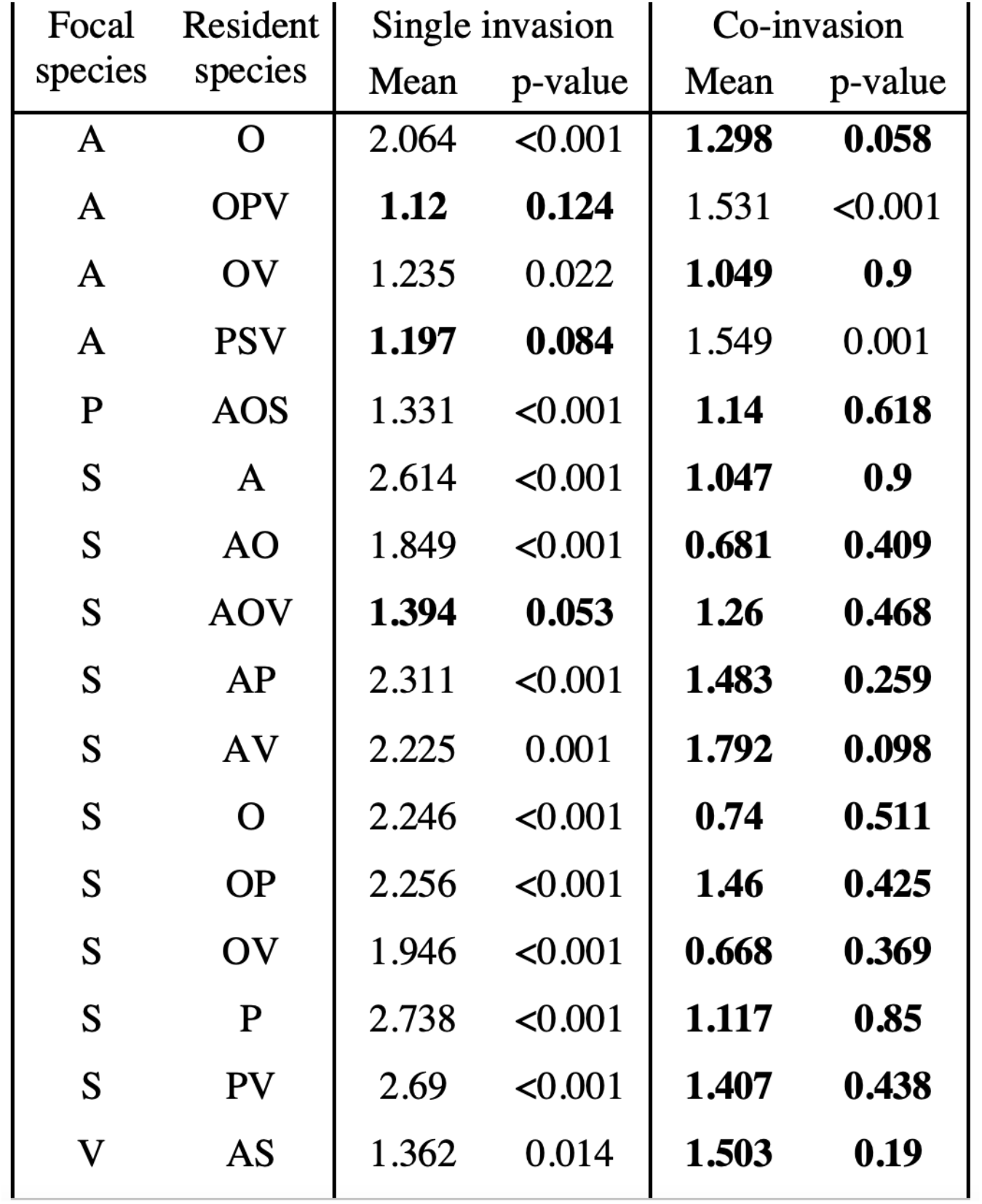
Mean estimates of relative invader fitness when focal species are invading into a community of different resident species as a single species or in the presence of co-invaders (remaining invading species not listed as either focal or residents). Listed are combinations in which at least one invasion across the two invasion types failed. P-values (corrected for multiple testing) indicate the difference of estimates to 1 with non-significant values (bold) indicating failed invasions. Letters dictate the first letter of each species name.

Comparatively, when species are invaded in the absence of co-invaders into each combination of residents, 67/70 combinations were stable (Table 1; Figure 1) (Castledine *et al*. 2020b). One instance of failed invasion of *Stenotrophomonas* into residents of *Achromobacter, Ochrobactrum* and *Variovorax* was predicted in both single and multispecies invasion assays. The other two failed invasions were otherwise successful in multi-species invasion. Overall, these results show estimates of community stability are robust across levels of resident and invader diversity for at least ⅘ species.

### Values of relative invader fitness are comparable

Next, we compared values of relative invader fitness between assays. If single species invasions perfectly predicted relative invader fitness with co-invaders, a 1-1 relationship would be evident. Although there was a significant positive relationship between mean estimates of relative invader fitness, the slope was 0.159 (F_1,69_ = 4.93, p = 0.03; species-specific intercepts, F_4,69_ = 16.77, p < 0.001; interaction between relative invader fitness for single invasions and species: F_4,65_ = 0.594, p = 0.669). For *Achromobacter* sp. (8/14), *Pseudomonas* sp. (11/14) and *Stenotrophomonas* sp. (14/14), estimates of relative invader fitness in single invasions overestimated fitness in co-invasion assays. This suggests the presence of co-invaders decreases the relative invader fitness of these species. In contrast, most estimates of relative invader fitness for *Ochrobactrum* sp. (12/14) and *Variovorax* sp. (10/14) were underestimated compared to when co-invaders were present. As such, for these two species, the presence of co-invaders resulted in greater estimates of relative invader fitness.

However, as mean estimates do not account for within treatment variation, we next compared relative invader fitness values within combinations of resident species. Here, there was a significant interaction between treatment (focal species and resident species combination) and invasion type (presence or absence of co-invaders) (F_69,700_ = 10.91, p < 0.001). Comparisons within treatments revealed no significant differences in relative invader fitness in the presence and absence of co-invaders in 42/70 comparisons. 11/28 significant comparisons were accounted for by *Stenotrophomonas*, the species which had multiple failed invasions as above. 9/28 significant comparisons were accounted for by *Ochrobactrum* where in 7, estimates were significantly greater with co-invaders present. *Achromobacter* accounted for 4, *Pseudomonas* and *Variovorax* 2 each of the remaining significant comparisons. These results suggest that for most species within this community, estimates of relative invader fitness are robust to the presence of co-invaders when accounting for within-treatment variation.

## Discussion

The invasion criterion has become a widely used metric of community stability, however its applicability when multiple species decrease to low densities has not been experimentally tested (Grainger *et al*. 2019). Here, we show that single species invasions can predict community stability in most ecological scenarios in which multiple co-invaders are present. Mean estimates of relative invader fitness were less comparable among species between invasion assays with some species RIF being increased or decreased by the presence of co-invaders. However, within treatments, most RIF values were non-significantly different in ⅗ species showing high levels of repeatability in the presence and absence of co-invaders. As such, our study suggests that the invasion criterion which typically focuses on single-species invasions can provide accurate estimates of community stability.

In our previous study, this model community was shown to be stable in 67/70 combinations of two to four species (and ⅘ invasion assays with five species) showing a high-level of stability (Castledine *et al*. 2020b). We repeated these invasion assays with the same combinations of resident species and invaded all non-residents from rare, resulting in co-invasions of two to four species into communities of three to one species respectively. Here, 56/70 combinations showed stability with each species capable of invading from rare. *Stenotrophomonas* was predominantly affected by the presence of co-invaders and accounted for 10/14 failed invasions while this species only had 1 failed invasion as a single invader. We do not know the mechanism by which *Stenotrophomonas* was excluded but this may be due to competitive exclusion from faster growing co-invaders and/or being of lower density in the ratio of co-invaders (lower propagule pressure compared to single invasion). Comparatively, for ⅘ species, community stability was accurately predicted in the presence and absence of co-invaders with most RIF estimates being significantly greater than 1.

Furthermore, values of RIF were broadly predictable between invasion assays. For all species, a positive relationship between RIF estimates in single- and co-invasions was shown indicating a degree of predictability between estimates. Additionally, when comparing values of RIF between invasion assays for each combination of resident species, estimates were non-significantly different in 42/70 comparisons. Where comparisons were significantly different, *Stenotrophomonas* and *Ochrobactrum* accounted for 20/28 with *Stenotrophomonas* having multiple failed invasions and *Ochrobactrum* showing generally higher RIF when co-invaders are present. Comparatively, in ⅗ species, most estimates were predictable between assays showing RIF was mostly robust to the presence of co-invaders. This level of predictability is surprising with assays varying in how many species are present and all species being able to impact the fitness of others (Castledine *et al*. 2020b).

That co-invaders do not affect RIF values for ⅗ species or stability estimates for ⅘ species suggests that community stability is predominantly maintained by species occupying distinct niches and being limited by intra rather than interspecific competition (Castledine *et al*. 2020b; Chesson 2018). There is selection across taxa, observed from microbes to birds and fish, to reduce competition between species and diversify into separate niches (ecological character displacement) (Germain *et al*. 2018; Pfennig & Pfennig 2010). The invasion criterion has been applied to show evolution has resulted in species diversifying into separate niches with evolved populations having negative frequency dependent fitness (Christie & McNickle 2023). In such communities, the invasion criterion working with single species invasions may accurately predict community stability and the relative performance of each species when recovering from rare. Validity of RIF estimates of species that depend on others for resources is a particular concern when multiple species decline as these can result in extinction cascades (Pires *et al*. 2020). For instance, loss of mutualistic corals and fish following temperature extremes has resulted in co-extinction (Strona *et al*. 2021). None of the species in this community were obligate mutualists which perhaps increased the repeatability of our estimates. Comparing RIF estimates under single- and co-invasion assays when species interact mutualistically would be a next-step in understanding the broad validity of the invasion criterion in communities dominated by different interaction types.

## Acknowledgements

This work was supported by NERC awards NE/V012347/1 and NE/S000771/1 awarded to AB. DP is funded by a NERC independent research fellowship (NE/W008890/1).

## References

Adler, P.B., HilleRisLambers, J., Kyriakidis, P.C., Guan, Q. & Levine, J.M. (2006). Climate variability has a stabilizing effect on the coexistence of prairie grasses. Proc. Natl. Acad. Sci. U.S.A., 103, 12793–12798.

Altieri, A.H., van Wesenbeeck, B.K., Bertness, M.D. & Silliman, B.R. (2010). Facilitation cascade drives positive relationship between native biodiversity and invasion success. Ecology, 91, 1269–1275.

Angert, A.L., Huxman, T.E., Chesson, P. & Venable, D.L. (2009). Functional tradeoffs determine species coexistence via the storage effect. Proc. Natl. Acad. Sci. U.S.A., 106, 11641–11645.

Benjamini, Y. & Hochberg, Y. (1995). Controlling the False Discovery Rate: A Practical and Powerful Approach to Multiple Testing. Journal of the Royal Statistical Society: Series B (Methodological), 57, 289–300.

Buche, L., Spaak, J.W., Jarillo, J. & De Laender, F. (2022). Niche differences, not fitness differences, explain predicted coexistence across ecological groups. Journal of Ecology, 110, 2785–2796.

Castledine, M., Padfield, D. & Buckling, A. (2020a). Experimental (co)evolution in a multispecies microbial community results in local maladaptation. Ecol Lett, 23, 1673–1681.

Castledine, M., Pennycook, J., Newbury, A., Lear, L., Erdos, Z., Lewis, R., et al. (2020b). haracterising a stable five-species microbial community for use in experimental evolution and ecology.

Chesson, P. (2000). Mechanisms of Maintenance of Species Diversity. Annu. Rev. Ecol. Syst., 31, 343–366.

Chesson, P. (2018). Updates on mechanisms of maintenance of species diversity. J Ecol, 106, 1773–1794.

Christie, M.R. & McNickle, G.G. (2023). Negative frequency dependent selection unites ecology and evolution. Ecology and Evolution, 13, e10327.

Coblentz, K.E. & DeLong, J.P. (2020). Predator-dependent functional responses alter the coexistence and indirect effects among prey that share a predator. Oikos, 129, 1404–1414.

Descamps-Julien, B. & Gonzalez, A. (2005). Stable Coexistence in a fluctuating environment: an experimental demonstration. Ecology, 86, 2815–2824.

Gendron, R.P. (1987). Models and Mechanisms of Frequency-Dependent Predation. The American Naturalist, 130, 603–623.

Germain, R.M., Williams, J.L., Schluter, D. & Angert, A.L. (2018). Moving Character Displacement beyond Characters Using Contemporary Coexistence Theory. Trends in Ecology & Evolution, 33, 74–84.

Godoy, O., Stouffer, D.B., Kraft, N.J.B. & Levine, J.M. (2017). Intransitivity is infrequent and fails to promote annual plant coexistence without pairwise niche differences. Ecology, 98, 1193–1200.

Grainger, T.N., Levine, J.M. & Gilbert, B. (2019). The Invasion Criterion: A Common Currency for Ecological Research. Trends in Ecology & Evolution, 34, 925–935.

Lenski, R.E., Rose, M.R., Simpson, S.C. & Tadler, S.C. (1991). Long-Term Experimental Evolution in Escherichia coli. I. Adaptation and Divergence During 2,000 Generations. Am Nat, 138, 1315–1341.

Lenth, R. (2018). Emmeans: Estimated marginal means, aka least-squares means. R Package Version.

Li, S., Tan, J., Yang, X., Ma, C. & Jiang, L. (2019). Niche and fitness differences determine invasion success and impact in laboratory bacterial communities. The ISME Journal, 13, 402–412.

Petraitis, P.S., Latham, R.E. & Niesenbaum, R.A. (1989). The Maintenance of Species Diversity by Disturbance. The Quarterly Review of Biology, 64, 393–418.

Pfennig, D.W. & Pfennig, K.S. (2010). Character Displacement and the Origins of Diversity. The American Naturalist, 176, S26–S44.

Pires, M.M., O’Donnell, J.L., Burkle, L.A., Díaz-Castelazo, C., Hembry, D.H., Yeakel, J.D., et al. (2020). The indirect paths to cascading effects of extinctions in mutualistic networks. Ecology, 101, e03080.

Rainey, P.B. & Travisano, M. (1998). Adaptive radiation in a heterogeneous environment. Nature, 394, 69–72.

Sala, O.E., Stuart Chapin, F., Iii Armesto, J.J., Berlow, E., Bloomfield, J., et al. (2000). Global Biodiversity Scenarios for the Year 2100. Science, 287, 1770–1774.

Schoener, T.W. (1974). Resource Partitioning in Ecological Communities: Research on how similar species divide resources helps reveal the natural regulation of species diversity. Science, 185, 27–39.

Schwartz, D.J., Langdon, A.E. & Dantas, G. (2020). Understanding the impact of antibiotic perturbation on the human microbiome. Genome Med, 12, 82.

Siepielski, A.M. & McPeek, M.A. (2010). On the evidence for species coexistence: a critique of the coexistence program. Ecology, 91, 3153–3164.

Singh, P. & Baruah, G. (2021). Higher order interactions and species coexistence. Theor Ecol, 14, 71–83.

Stachowicz, J.J. (2001). Mutualism, Facilitation, and the Structure of Ecological Communities: Positive interactions play a critical, but underappreciated, role in ecological communities by reducing physical or biotic stresses in existing habitats and by creating new habitats on which many species depend. BioScience, 51, 235.

Stomp, M., Huisman, J., De Jongh, F., Veraart, A.J., Gerla, D., Rijkeboer, M., et al. (2004). Adaptive divergence in pigment composition promotes phytoplankton biodiversity. Nature, 432, 104–107.

Strona, G., Beck, P.S.A., Cabeza, M., Fattorini, S., Guilhaumon, F., Micheli, F., et al. (2021). Ecological dependencies make remote reef fish communities most vulnerable to coral loss. Nat Commun, 12, 7282.

Team, R.C. (2013). R: a language and environment for statistical computing.

Wickham, H. (2016). ggplot2: elegant graphics for data analysis. Springer, New York.

Winter, C., Bouvier, T., Weinbauer, M.G. & Thingstad, T.F. (2010). Trade-Offs between Competition and Defense Specialists among Unicellular Planktonic Organisms: the “Killing the Winner” Hypothesis Revisited. Microbiol. Mol. Biol. Rev., 74, 42–57.

